# SARS-CoV2 genome analysis of Indian isolates and molecular modelling of D614G mutated spike protein with TMPRSS2 depicted its enhanced interaction and virus infectivity

**DOI:** 10.1101/2020.07.23.217430

**Authors:** Sunil Raghav, Arup Ghosh, Jyotirmayee Turuk, Sugandh Kumar, Atimukta Jha, Swati Madhulika, Manasi Priyadarshini, Viplov K. Biswas, P. Sushree Shyamli, Bharati Singh, Neha Singh, Deepika Singh, Ankita Datey, Avula Kiran, Shuchi Smita, Jyotsnamayee Sabat, Debdutta Bhattacharya, Rupesh Dash, Shantibhushan Senapati, Tushar K. Beuria, Rajeeb Swain, Soma Chattopadhyay, Gulam Hussain Syed, Anshuman Dixit, Punit Prasad, Odisha COVID-19 study group, ILS COVID-19 team, Sanghamitra Pati, Ajay Parida

**Author notes:** Equal contribution first author (Sunil Raghav, Arup Ghosh). Corresponding author **Correspondence to:** Dr. Ajay Parida, Dr. Sunil Raghav. This work is done under the umbrella of DBT’s PAN-India 1000 SARS-CoV2 RNA genome sequencing consortium. **Odisha COVID-19 study group** (Author names are arranged in alphabetical manner): Arvind Kumar Singh, Baijayantimala Mishra, Banajini Parida, Binod Kumar Patro, D. P. Dogra, Dasarathi Das, Deepa Prasad, Dhaneswari Jena, Dharitri Mohapatra, Dinesh Prasad Sahu, Durga Madhab Satapathy, Durgesh Prasad Sahoo, Jayanta Panda, Jaya Singh Khatri, Kaushik Mishra, Manoranjan Satpathy, Nirupama Chaini, Roma Rattan, Sadhu Panda, Sangeeta Das, Somen Kumar Pradhan, Srikanta Kanungo, Sriprasad Mohanty, Subrata Kumar Palo. **ILS COVID-19 group (Author names are arranged in alphabetical manner)**: Aditi Chatterjee, Ajit Singh, Amol Suryawanshi, Amrita Ray, Ankita Datey, Aliva Minz, Auromira Khuntia, Debyashreeta Barik, Deepak Singh, Deepika Singh, Dileep Vasudevan, Eshna Laha, Hiren G. Dodia, Kautilya Jena, Kaushik Sen, Manashi Priyadarshini, Omprakash Shriwas, P. M. Vaishali, Parej Nath, Prabhudutta Mamidi, Priyanka Mohapatra, Rahul Das, Reena Yadav, Satyaranjan Sahoo, Saikat De, Sanchari Chatterjee, Sandhya Suranjika, Shamima Ansari, Shiva Pradhan, Sivaram Krishna, Sneha Dutta, Soumyajit Gosh, Subhabrata Barik, Tsheten Sheroa.

## Abstract

COVID-19 that emerged as a global pandemic is caused by SARS-CoV-2 virus. The virus genome analysis during disease spread reveals about its evolution and transmission. We did whole genome sequencing of 225 clinical strains from the state of Odisha in eastern India using ARTIC protocol-based amplicon sequencing. Phylogenetic analysis identified the presence of all five reported clades 19A, 19B, 20A, 20B and 20C in the population. The analyses revealed two major routes for the introduction of the disease in India i.e. Europe and South-east Asia followed by local transmission. Interestingly, 19B clade was found to be much more prevalent in our sequenced genomes (17%) as compared to other genomes reported so far from India. The haplogroup analysis for clades showed evolution of 19A and 19B in parallel whereas the 20B and 20C appeared to evolve from 20A. Majority of the 19A and 19B clades were present in cases that migrated from Gujarat state in India suggesting it to be one of the major initial points of disease transmission in India during month of March and April. We found that with the time 20A and 20B clades evolved drastically that originated from central Europe. At the same time, it has been observed that 20A and 20B clades depicted selection of four common mutations i.e. 241 C>T (5’UTR), P323L in RdRP, F942F in NSP3 and D614G in the spike protein. We found an increase in the concordance of G614 mutation evolution with the viral load in clinical samples as evident from decreased Ct value of spike and Orf1ab gene in qPCR. Molecular modelling and docking analysis identified that D614G mutation enhanced interaction of spike with TMPRSS2 protease, which could impact the shedding of S1 domain and infectivity of the virus in host cells.

## Introduction

Coronavirus disease 19 (COVID-19) pandemic is caused by severe acute respiratory syndrome coronavirus 2 (SARS-CoV-2), a betacoronavirus belonging to the coronaviridae family. The first occurrence of this novel coronavirus was observed in Wuhan, China in late December, 2019 which later spread globally via human-to-human contact transmission [1]. According to the COVID-19 dashboard by Johns Hopkins University the virus has spread to more than 180 countries with 14.1 million total confirmed cases and 600,000 deaths worldwide [2].The first occurrence of coronavirus case in India was observed in mid-January but the number of cases started to increase from the first week of March. According to official sources the number of cases in India has reached 1.2 million with 29,000 deceased. Genomic studies performed to understand the origin of SARS-CoV2, a positive single stranded RNA virus, unravelled that it has a zoonotic origin and is transmitted to humans from bats via malayan pangolins [3] .The nucleotide sequence of SARS-CoV2 is ~79% similar to SARS-CoV1 and about 50% with Middle East Respiratory Syndrome coronavirus (MERS-CoV) [1]. Approximately 30 kb genome of SRAS-CoV2 features a cap structure in 5’ and 3’ poly (A) like other members of the coronavirus family. Major portion of the genome is covered by two ORFs (ORF1a and ORF1b) which codes for 15 non-structural proteins including crucial proteins required for viral replication like viral proteases nsp3, nsp5 and nsp12 also known as RNA-dependent RNA polymerase (RdRP) [4]. The gRNA also codes for structural proteins like spike protein(s), nucleocapsid protein (N), membrane protein (M) and envelope protein (E) required for packaging of the virus and at least accessory proteins but their ORFs are still not experimentally validated [4].

The entry of viral particles in the human body occurs through binding with angiotensin I converting enzyme 2 (*hACE2)* receptor present on lung epithelial cells [5]. The predominant infection site is respiratory tract due to route of infection. Other than lungs it is known to infect other organs of the body such as kidney, liver and intestine as ACE2 expression is found to be quite high. After the viral entry into host cells, to initiate viral replication first negative-sense RNA intermediates are synthesized by RdRP activity, these templates are then utilized for synthesis of gRNA and sub-genomic RNAs (sgRNAs) [4]. Because of the low-fidelity of RdRP, mutations are incorporated with high frequency in gRNA [6] and often such mutations are known to increase the pathogenicity and fitness of the virus [7],[8].

Recently a predominant mutation i.e. D614G in spike protein has been identified in virus strains sequenced from European population [8].There are several reports depicting that D614G mutation in spike protein is associated with enhanced infectivity and spread of the virus owing to increased interaction with ACE2 receptor present on host cells [8]. It has also been indicated that this mutation is present on the S2 domain of spike protein that is important for cleavage by TMPRSS2 enzyme for cleavage of S1 for facilitating the fusion of the viral spike with the host cell membrane [8, 9]. It is indeed interesting to understand further at molecular level, the changes in protein structure induced by this mutation and its functional association with the infectivity and disease severity.

At the same time, it has been reported that D614G mutation co-occurs with 3 more mutation i.e. 241 in UTR, 3307 and 14408 [8]. The functional significance of these co-occurring mutations in evolution and selectivity of virus is interesting to understand.

In the present study we have sequenced 225 COVID-19 isolates from patient samples from the state of Odisha those migrated from 13 most affected Indian states as a part of DBT’s PAN-INDIA 1000 SARS-CoV2 RNA genome sequencing consortium, using amplicon sequencing based methodology. The travel history of these patients was collected along with their symptomatic and asymptomatic behaviour. We performed phylogenetic analysis from sequenced data to understand the genetic diversity and evolution of SARS-CoV2 in the Indian subcontinent. From the sequenced isolates we have identified 247 single nucleotide variants most of them are observed in ORF1ab, spike and nucleocapsid protein coding region. Moreover, we have analysed the D614G mutations in the samples to obtain information on its evolution in Indian population. Protein-modelling analysis of D614G mutation was carried out to identify the impact on structural changes at protein levels. We further performed protein-protein docking simulation to predict the impact of D614G mutation on the interaction between of wild-type and mutated spike protein with TMPRSS2 enzyme to assess its impact on binding and perhaps viral infectivity.

## Results

### Demographics, clinical status and travel history

The average age of the 225 subjects were 30.98 ± 11.79 years with age range 1-75 years and median age of 30 years. The overall gender ratio of male: female was 201: 24 with median age of male subjects 31.75 years and 24.5 years for female subjects (Supplementary Figure 1A, 1B). Almost every female subject in this study reported no strong symptoms during sample collection whereas in case of the male subjects we found that the numbers of symptomatic cases were almost same with the number of asymptomatic cases (Supplementary Figure 1C). For the samples in our study, we didn’t observe any fatality in the subject group to our knowledge.

Majority of the subjects (89%) disclosed their travel history during the sample collection procedure. We found that the majority of the subjects included in the study were found to migrate from Gujarat, West Bengal, Tamil Nadu, Maharashtra, Delhi, Kerala and Andhra Pradesh state in India (Supplementary Figure 1D). As most of the subjects did not had any direct foreign travel (n = 1) history the primary source of infection is local contacts in their workplaces and this helped us to understand the COVID-19 strain diversity in the most affected states of India (Supplementary Figure 1D).

### Mutation analysis

After filtering out low quality samples with <5% N’s in the assembly and at least 10x average coverage we observed a total of 247 single nucleotide variants from 202 SARS-CoV2 isolates. Out of these variants 156 variants observed only in single isolates, 25 variants were classified as common variants with occurrence in more than 5% isolates and 19 variants as rare with 2-5% occurrence in all samples (Supplementary Table 1). Among the common variants the most frequent mutations are 23403 A>G (D614G, S gene), 241 C>T (5’ UTR), 14408 C>T (P4715L, RdRP gene), 3037 C>T (F942F, NSP3), 28881 G>A (R203K, N gene), 28882 G>A (R203R, N gene), 28883 G>C (G204R, N gene), 28144 T>C (L84S, ORF8) with presence in more than 15% of all of our sequenced samples (Supplementary Table 1). Plotting mutation diversity (>2%) in a clade wise manner we observed distinct mutation signatures in different clades. The isolates that were grouped in 19A clade depicted prevalence of mostly ORF1ab mutations with one distinct N gene C>T mutation at 28311 position (Figure 2C). In clade 19B samples we observed a very distinct ORF8 T>C mutation at 28144 position, two N gene mutations at position 28326 and 28878 with some ORF1ab mutation in lower frequency (Figure 2C). Clade 20A and 20B has almost similar mutation profile with major mutated positions 241 C>T mutation in leader sequence, ORF1ab 3037 C>T, ORF1ab 14408 C>T (RdRP) and S gene 23403 A>G mutations. The very distinct characteristic that we observed for 20B clade is three consecutive N gene mutations at position 28881, 28882 and 28883 (Figure 2C). Out of these three mutations two are missense mutations resulting in change of protein sequence. In clade 20A we observed two mutations at ORF3a and M protein coding gene at position 25563 G>T, 26735 C>T but in less frequencies in comparison with other mutated sites (Figure 2C). Overall, to conclude we summarised all the mutated sites present in all the samples and observed ORF1ab is the most mutated region followed by N gene, S gene and ORF8 in SARS-CoV2 samples from India (Figure 2A). Among the ORF1ab mutation 4715 P>L change in nsp12 also known as RdRP (the viral RNA dependent RNA polymerase) was the most prevalent one followed by a synonymous change (F924F) in nsp3 protein (Figure 2B).

**Figure 1:**
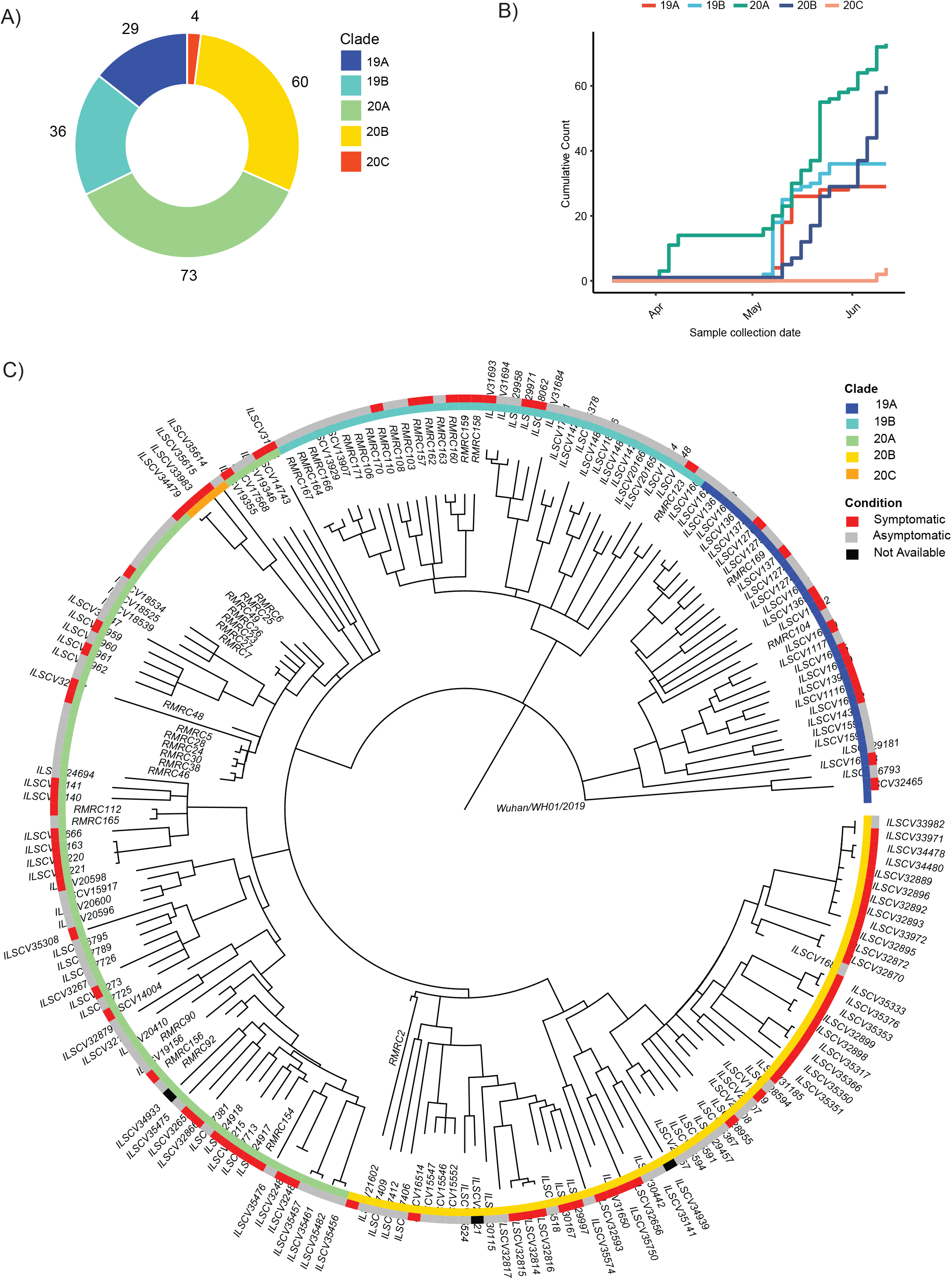
Phylogenetic analysis of the SARS-Cov genomes and their distribution into different Nextstrain defined new clades. (A) A donut chart representing the sequenced sample (n=202) distribution across the clades (clade nomenclature obtained using Nextstrain). (B) Cumulative count of clades plotted against sample collection date showing abundance of clades with time. (C) Time-tree of the sequenced samples (n = 202) generated using Nextstrain Augur overlaid with clinical status (Condition) of the patient during sample collection and clade information.

**Figure 2:**
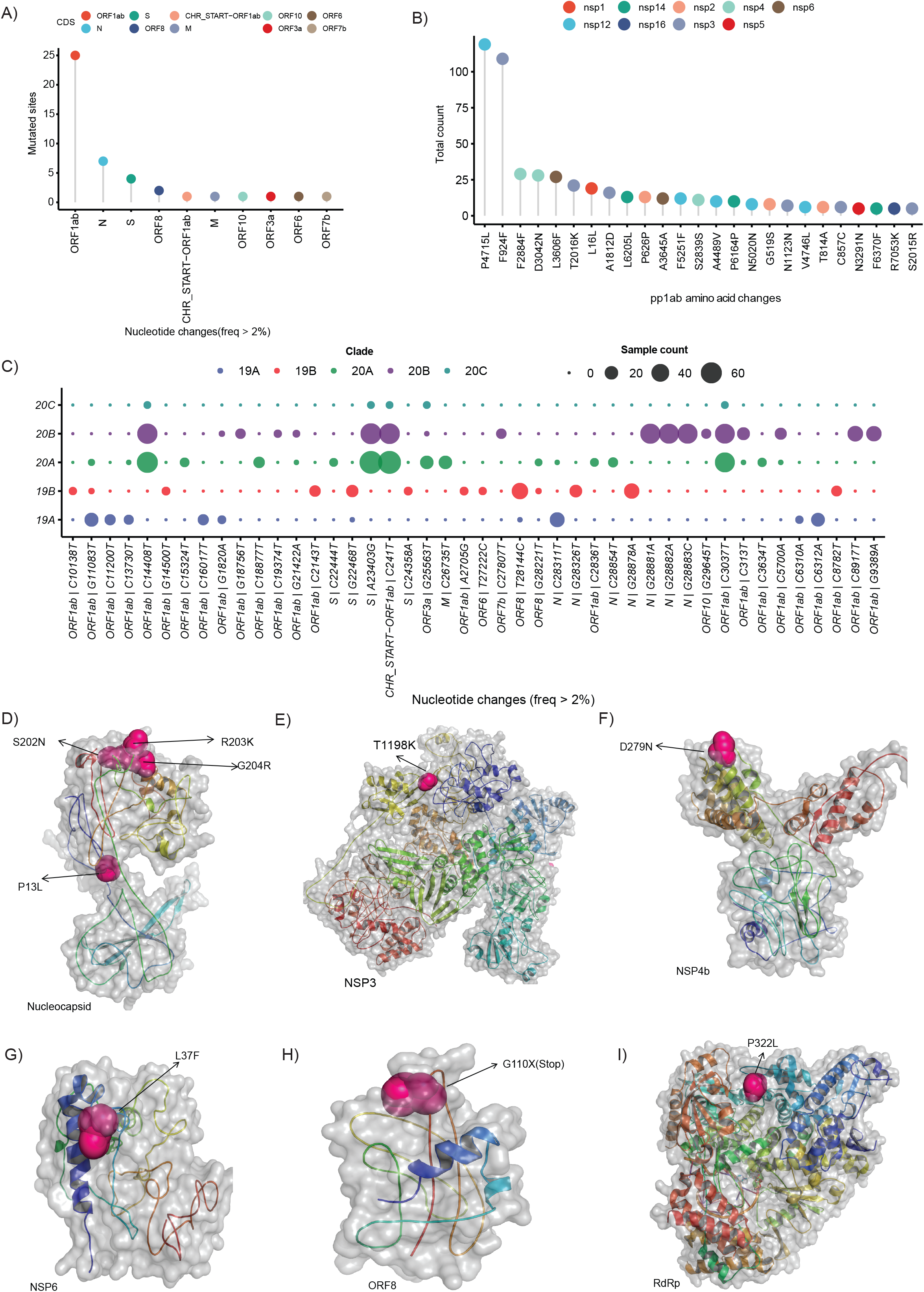
SARS-CoV2 clade distribution and their prevalent mutation profiles. (A) Dot plot representing number of single nucleotide mutation (occurred in more than 2% of the samples) present in different genomic segments of SARS-CoV-2 genome. (B) The ORF1ab region codes for a polypeptide is later cleaved to several mature peptides. The dot plot represents the amino acid changes (location of amino acid acids as per location in polypeptide sequence) in the mature peptides of ORF1ab (C) Clade wise occurrence of nucleotide mutations with presence in more than 2% of sequenced samples (n = 202). Colour of the dots represents the clade and size of the dots represents number of the samples showing presence of the single nucleotide variant. (D-I) The mutation sites on the modelled structures of the SARS-CoV2 proteins. The mutation site(s) of the NSP3, NSP4b, NSP6, RdRP and nucleocapsid proteins are marked as sphere while the rest of the structure is shown in cartoon representation.

### Haplotype network analysis

First we created a median-joining haplotype network to look for transmission within India we observed two major branches (Supplementary Figure 2A). When we coloured the sequences with clade information the branched represented 19A and 20A which later furcate into 19B, 20B and 20C (Supplementary Figure 2A). When we overplayed migration information in the network we observed majority of 19A and 19B isolates migrated from Gujarat (Figure 3A). Migration of isolates identified in newly prevalent clades occurred from southern path of India (Figure 3A). To understand the source transmission of SARS-CoV-2 infection in India, we constructed haplotype networks using sequencing data combined with genome sequences from the countries (China 15, Germany 23, Italy 25, Saudi Arabia 23, Singapore 14 and South Korea) from where the migration of individuals has occurred to India. From the haplotype network we observed distinct clusters of genome sequences that were grouped in 4 major nodes. A large group of sequences clustered in two major haplotype clusters one with genome sequences from China, Singapore, South Korea and the other one with Italy, Saudi Arabia and Germany with 2-4 nucleotide substitutions (Figure 3B). From the collection date we observed that 20A clade which is prevalent in Europe, became abundant in Odisha from April representing the top cluster, while the bottom cluster represents samples belonging to clade 19A, 19B having a common origin in South-east Asia (Figure 3B).

**Figure 3:**
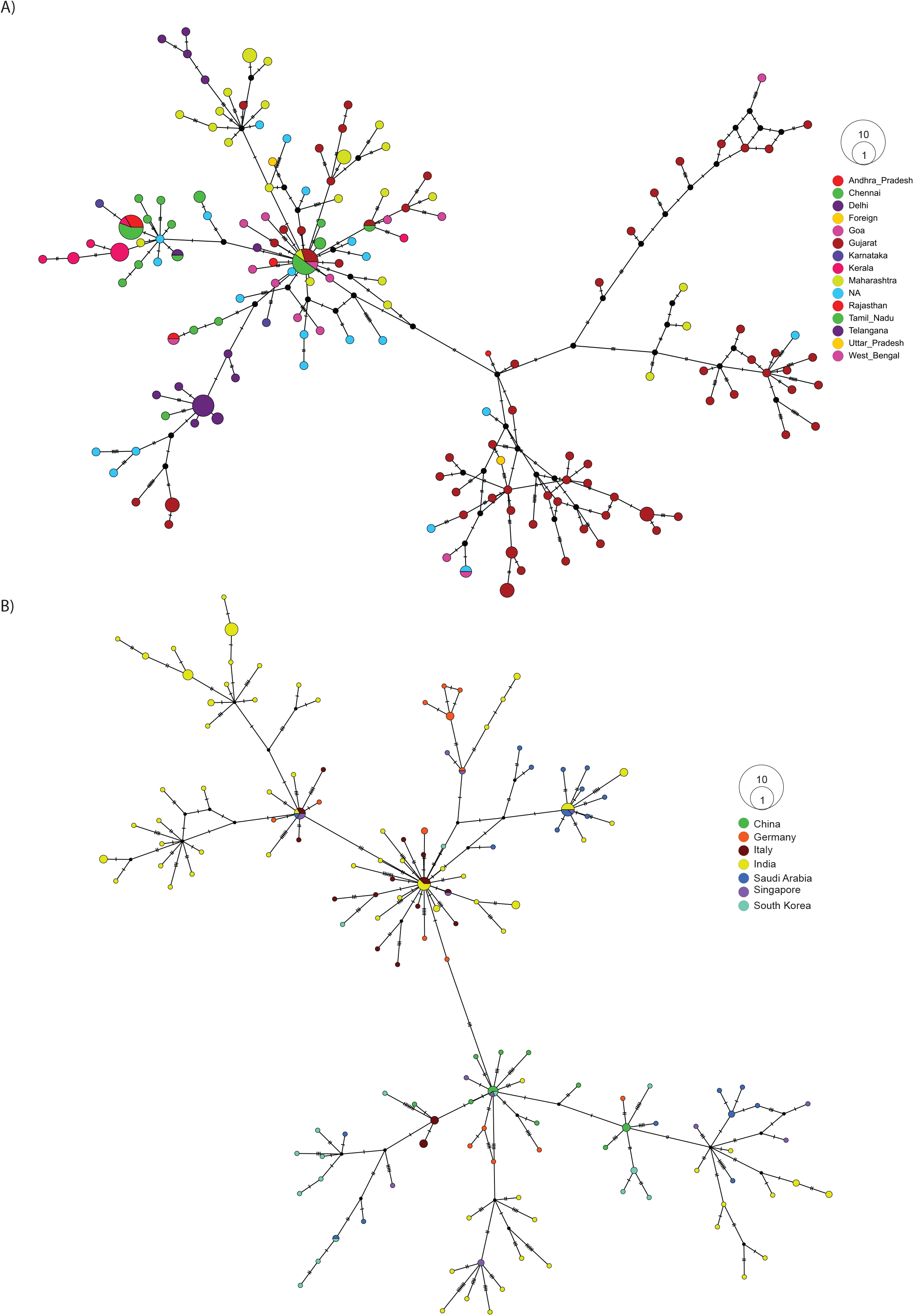
Haplotype network analysis of SARS-CoV-2 sequences. (A) Haplotype network of 202 SARS-CoV-2 whole genome sequences from our dataset coloured by their respective place of migration. (B) Haplotype network of 100 high coverage SARS-CoV2 genomes obtained from GISAID (China 15, Germany 23, Italy 25, Saudi Arabia 23, Singapore 14, South Korea) combined with 187 samples sequenced from Odisha with less than <5% N’s present in consensus sequence.

### Phylogenetic analysis

The phylogenetic analysis was carried out with 202 high quality SARS-CoV2 sequences that revealed presence of four major clades i.e. 19A (n = 39), 19B (n = 36), 20A (n = 73), 20B (n = 60) and one minor clade 20C (n = 4) (Figure 1 A). To understand the abundance of clades with time, we plotted cumulative counts of clades against the sample collection date and observed the introduction of clade 20A occurred in April, 2020 with parallel emergence of 19A, 19B and 20B in May (Figure 1B). After May no new occurrence of clade 19A or 19B were observed in our data but in mid-June we started to observe emergence of 20C clade (Figure 1B). Further to understand the transmission of the virus in our subjects, we performed phylogenetic analysis of our dataset with 1042 high coverage Indian SARS-CoV2 whole genome sequences (N <1%) obtained from GISAID on 12th July 2020 (Supplementary Figure 1F). From the transmission map generated using Nextstrain, we observed that most of the sequences in 20A clades share sequence similarity with samples from Gujarat, which has very high prevalence of this clade (Supplementary Figure 1E). The 19A clade is very prevalent in Delhi and Telangana region and Tamil Nadu but the 20B clade is only prevalent in the southern parts of India (Supplementary Figure 1E). Both 20A and 20B clade predominantly contain a mutation in Spike protein coding sequence 23403A>G and the mutation is traced back to the West European region in late January [8]. The missense mutation in spike protein coding gene causes a change in 614 D>G position of Spike protein and reported to increase the shedding of S1 subunit of the protein which leads to the increased infectivity [8]. To understand the prevalence and effect of G614 in our sequencing dataset we plotted week wise count (based on collection date) of D614 and G614 and observed that the occurrence of the mutation has been observed in early march (12^th^ week) and the cumulative frequency increased over the time (Figure 4A). Every sample we sequenced as a part of the study was also checked for the levels of ORF1 and S gene using quantitative PCR. The Ct obtained is also a direct indicator of viral load (lesser the Ct, higher the viral load) in the individual. When we plotted the Ct values of all sequenced samples, we observed that except week of 21 the Ct values of the isolates having G614 mutation is less in comparison to isolates having D614 (Figure 4B, 4C). We also had Ct values (ORF1, S gene) of 637 positive isolates available as a partner institute of the COVID19 surveillance program in Odisha, India. Plotting the data against the date of sample collection for testing, we observed a sharp and significant (p<0.05) decline (median ~5 Ct change) in the Ct values (Figure 4E) from April, 2020 to May 2020 and there was median ~1 Ct change of S gene as well as ORF1ab (Figure 4D, 4E, 4F) from month of May to June 2020. In the case of ORF1 expression we also observed a sharp change between April and May but there is almost no change between May to June 2020 (Supplementary Figure 2A). When we overlaid the clinical status information (Asymptomatic *vs* symptomatic), we observed that isolates from clade 20A, 20B and 20C depicted higher number of symptomatic patients (Figure 1C). Although without the information about predisposition of complication in patients it’s hard to establish an association between mutations and the clinical manifestations.

**Figure 4:**
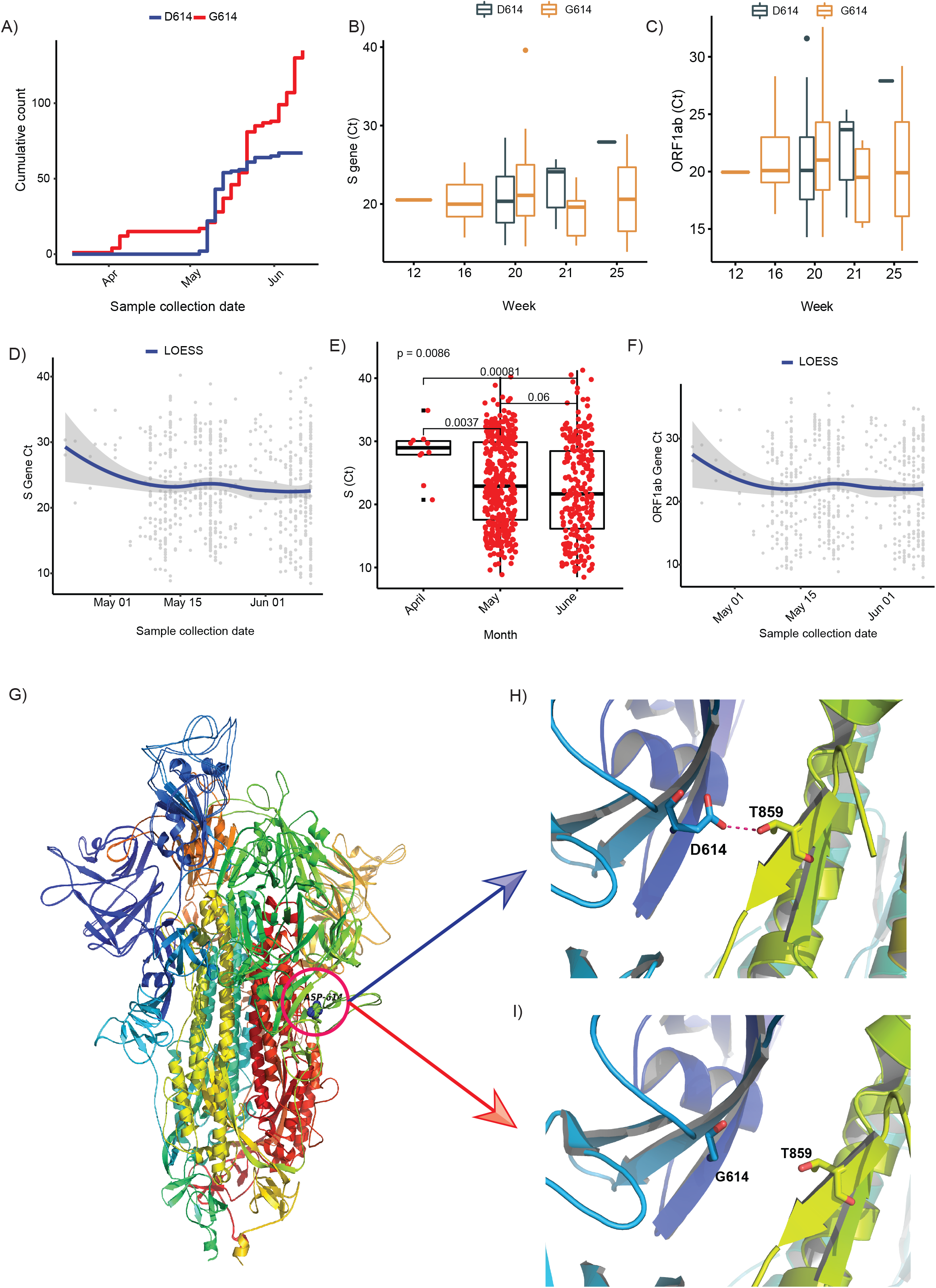
D614G in Spike gene increases infectivity potrayed by Ct values as a surrogate for viral load. (A) Cumulative count of the occurrence of D and G in 614 position of Spike protein in sequenced isolates (n=202). (B-C) Ct value distribution of S gene and ORF1ab for the sequenced isolates (n=202). (D-F) Ct value distribution of S gene and ORF1ab in all the positive samples tested at Institute of Life Sciences until June 17,2020. (G-I) The superimposed 3D-structures G614 mutant and wild type Spike protein. (G) The mutant site is highlighted with a circle at 614 position. (H) The hydrogen bond (D614-T859) shown as dotted line between Spike S1 and S2 domain in wild type. (I) The hydrogen bond is lost as a result of D614G mutation.

### Protein structure analysis

To understand the impact of identified prominent missense mutations (present on >10% of the samples) on the annotated viral proteins in our sequenced population, we used protein crystal structures and protein models as crystal structures were not available. The PROCHECK [10] results showed that the generated models have acceptable stereochemistry. First, we looked into the location of the mutated amino acid with respect to its functional domains as any change in the functional domain has high probability to perturb the protein function. The sites of mutation on the modelled protein structures (Nucleocapsid, NPS3, NSP4b, NSP6, ORF8 and RdRP) are marked to indicate their location (Figure 2D-2F). As reported in earlier publications, we also noticed D614G as highly prevalent mutation in highly transmitted and evolved strains belonging to clades 20A and 20B in Indian scenario, therefore we carried out molecular modelling analysis. It was observed that the wild-type (D614) and mutated form (G614) of Spike protein showed a slightly different arrangement of structural elements near the mutation site, which is present in hinge region linking S1 and S2 domain i.e. S1 furin cleavage site (Figure 4G). The S2 domain of the Spike protein is reported to interact with TMPRSS2 protease necessary for shedding of S1 domain for viral entry inside the host cells by facilitating the merging of virus with the host cell membrane. The proteolytic cleavage of spike protein by TMPRSS2 results in the formation of two fragments S1/S2, which is one of the key steps of the virus infection in host cells. The Spike protein contains a hydrogen bond between T859-D614 keeping S1 and S2 domains together (Figure 4H). The mutation of Asp to Gly at 614 eliminates this hydrogen bonding (Figure 4I) which enables a more favourable orientation of Q613 that may facilitate cleavage by TMPRSS2 by perturbing its affinity with S2 domain. It has also been proposed that D614 forms an intra-salt bridge with R646 which makes the conformation unfavourable for S1 association with S2 domain [9]. Interestingly, the protein docking analysis suggested better hydrogen bonding interactions between the spike protein cleavage sites (Arg685, Ser686) with the catalytic triad of TMPRSS2 in mutant condition as compared to wild-type. In the case of mutant Spike protein, the Arg682 and residues at primary cleavage site of Spike protein (Arg685 and Ser686) formed six hydrogen bonding interactions with Glu299, Lys300, Asp338, Gln438 residues of TMPRSS2 (Figure 5A, 5B). Whereas in the D614 wild-type form there were five hydrogen bonds observed between the cleavage site of Spike protein S2 domain and TMPRSS2 (Figure 5C, 5D). The binding energy was observed to be better for the G614 mutant (−143.03 kcal/mol) as compared to that of the wild type (−113.67 kcal/mol) indicating better binding of TMPRSS2 with the mutant Spike protein (Figure 5E).

**Figure 5:**
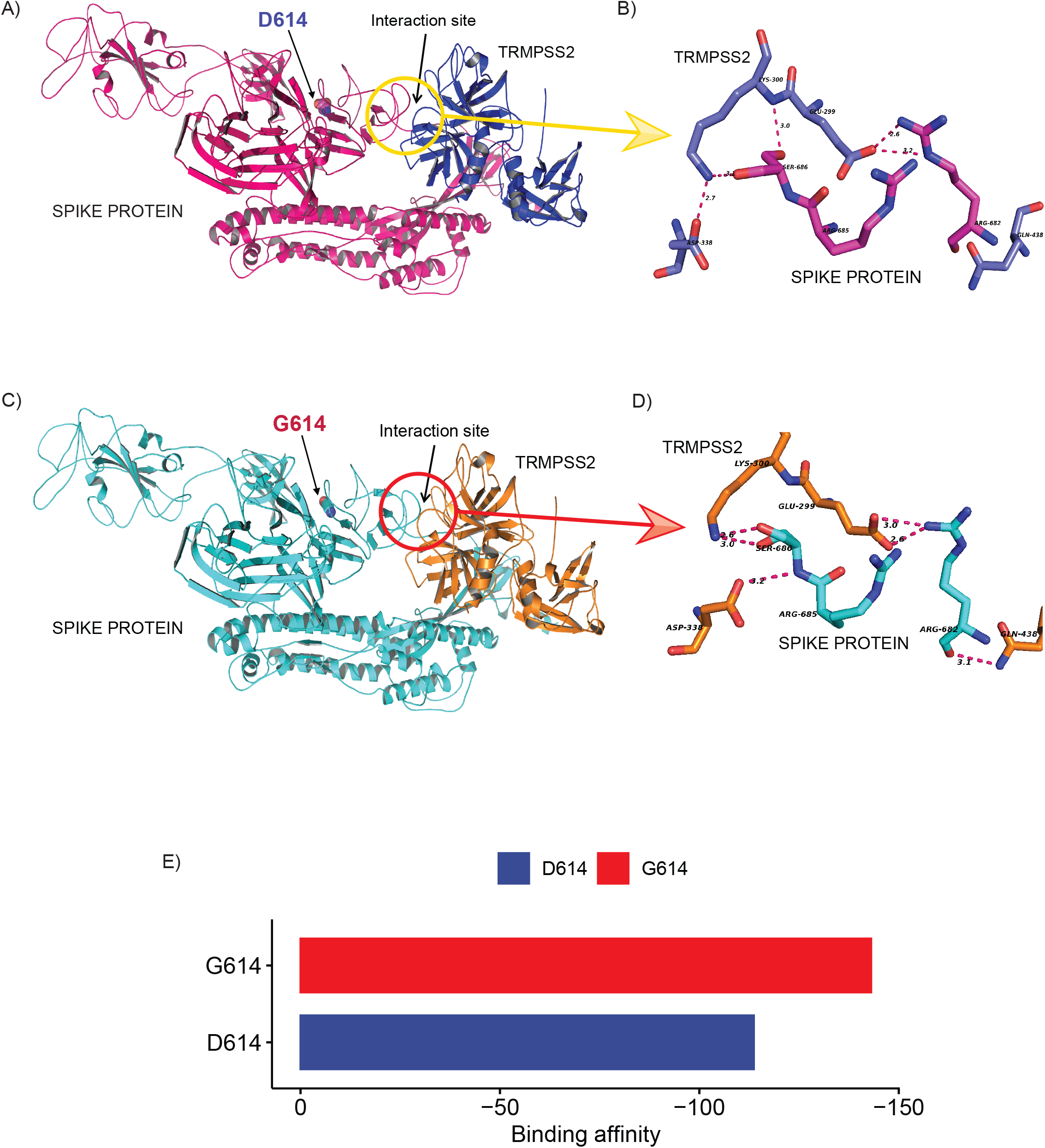
D614G change in Spike protein enhanced TMPRSS2 protease interaction that might be responsible for increased virus infectivity. (A-D). The docking study of TMPRSS2 with the wild-type (D614) and mutant (G614) Spike protein. The interaction site and the mutation position (614) is marked with an arrow. The hydrogen bond interactions are shown in pink dotted lines with distance marked in Å, (A) The overview of the docking site location on WT Spike protein, (B) The interactions between TMPRSS2 and wild type, (C) The overview of the docking site location on WT Spike protein, (D) The interactions between TMPRSS2 and mutant Spike protein, and (E) The average binding energy (kcal/mol) values for the top poses selected from five different clusters.

## Discussion

The pattern of COVID-19 pandemic spread in India is quite different from all over the world in term of slow transmission and lower mortality rates in the begining. Therefore, it was extremely important to understand the transmission dynamics of SARS-CoV2 and its evolution during different phases of disease in India. The mutations that accumulate in the virus genome with time acts as a molecular clock that can provide insight into emergence and evolution of the virus. These analyses could be helpful to prevent or control the transmission of the virus. To track the COVID-19 outbreak and understand the genomic clades of SARS-CoV2 prevalent in India as compared to rest of the world, we performed the genome sequencing of isolates collected from the oropharyngeal and nasopharyngeal swabs of individuals migrated from different regions of India to the state of Odisha after the reported incidences of COVID-19 pandemic in India.

For SARS-CoV2 genome sequencing, we did amplicon-based sequencing for 225 SARS-CoV2 genomes to capture the major clade diversity in India. We selected migrant groups from different parts of the country so that diversity of COVID-19 virus prevalent in the country could be captured in the phylogenetic analysis, to understand disease transmission and virus evolution with time. The sequenced samples represent migratory populations from North, East, West and Southern parts of India. Our analysis depicted that all the genomes analysed in our study were grouped into 4 major clades 19A, 19B, 20A and 20B according to the new Nextstrain clade nomenclature. The clade 19A is the Wuhan clade from China. Interestingly, we were able to capture occurrence of a rare clade with very less occurrence in India i.e. 19B with 17% (n=36) in our samples. This clade was found to be prevalent in East and South-east Asia during early outbreak of the pathogen. This confirmed that the foundation of the source of early outbreak infection in India also came from South-east Asian countries. The migration information for these early collected samples were not well defined therefore, it was difficult for us to exactly pinpoint from which South-east Asian country the transmission of 19B started. The mutation analysis depicted that both 19A and 19B clades almost evolved in parallel as prevalent mutations in both the clades are highly variable. On the other side, the phylogenetic analysis showed that clades 20A and 20B evolved quite rapidly in the Indian population and are major source of disease transmission in the country. Whereas 20C strain is rarely detected and appeared to be less adapted or somehow contracted by contact tracing at early stages of infection. The haplotype network construction also pointed to the later strains belonging to 20A, 20B clades originated from Western Europe and transmitted directly or via Saudi Arabia are mostly prevalent in the southern and western part of India. In the clade we also observed a very less frequent clade 20C only prevalent in a handful of the Middle-eastern countries makes up a marginal portion (n=4) of our sequenced samples. This suggests the requirement of constant monitoring of COVID19 isolates with sequencing technology to understand the source of infection and design prevention mechanisms like strategic lookdowns and region specific travel restrictions.

The fitness of the virus strain and its transmission depends on the adaptive mutations that it acquires with time. We found that four common variants i.e. 241 C>T in the UTR region, 3037 C>T in NSP3 gene, 14408 C>T in the NSP12 and 23403 A>G in S gene co-evolved mostly in the 20A and 20B clade. As 20A, 20B clade frequency increased with time in the population, which indicates that these strains have some selective advantage with time for increased transmission. It has been reported that the leader sequence present in the UTR region of positive strand RNA viruses like SARS-CoV is important for the replication and strand switching to generate negative strands [4]. This mutation in leader sequence is 20 nucleotides upstream of translation start site of ORF1ab gene. Therefore, it might be providing an advantage to virus in preventing stem loop generation required during strand switching by RdRP, which needs further experimental evaluation. At the same time, several reports documented that 23403 A>G (D614G) missense mutation in the Spike protein enhanced the infectivity rate of the virus. One of the reports showed that the shedding of S1 domain of Spike protein changes due to change in the hydrogen bonding between S1-S2 domain. We observed in our protein modelling analysis that, TMPRSS2 binding to Spike protein is enhanced by this mutation of Aspartic acid to Glycine (D614G) as it resulted in increased hydrogen bonding interactions. This change enhances the interaction of TMPRSS2 with the S2 domain, which is important for the cleavage of the S1 domain and virus entry into cells by facilitating its entry into the host cells. The overall primary in-silico docking study showed that mutant Spike protein has a greater number of hydrogen bonds with TMPRSS2 at the cleavage site as compared to the wild type resulting in better docking energy. Overall analysis indicates that the breakage of a hydrogen bond as a result of the mutation may facilitate greater cleavage of the mutant spike protein as compared to the wild type. All the sequenced genomes were submitted in the GISAID database.

## Materials & Methods

### Sample collection

All hospitalized and quarantined patients (March, 2020 to June, 2020) based on their clinical symptoms (fever or respiratory symptoms) or travel history, were preliminarily involved in this study. We received throat swabs in viral transport media (VTM) samples of these patients used for SARS-CoV-2 detection. Patients absent of or with negative SARS-CoV-2 test results were excluded from this study based on Ct values obtained by qPCR of isolated RNA. All patients involved in this study were residents of Odisha, India, during the outbreak period of COVID-19.

### Viral load detection

RNA isolation for all the 248 human subjects were performed using QIAamp Viral RNA Mini Kit (Qiagen, Cat. No. 52906). The isolated RNA was subjected to qPCR for determining viral load by Ct values. For qPCR we performed one-step multiplex real time PCR using TaqPath™ 1-Step Multiplex Master Mix (ThermoFisher Scientific, Cat. No. A28526) targeting 3 different gene-specific primer and probe sets - envelope glycoprotein spike (S), nucleocapsid (N), and open reading frame 1 (ORF1).

### Viral DNA library preparation & sequencing

We prepared a viral DNA library for viral genome sequencing using QIAseq FX DNA Library Kit (Qiagen, Cat. no. 180475) as instructed by the manufacturer’s manual and the library was subsequently sequenced using Illumina platform. The adapter sequence used for each sample was compatible with Illumina sequencing instrument with 96-sample configurations (Qiaseq unique dual Y-adapter kit). The average insert length was in the 250-500bp range. Prepared libraries were then pooled as a batch of 96 samples and sequenced using Illumina NextSeq 550 platform in 150 × 2 layout.

### Raw data pre-processing

Quality of the sequenced files were checked using FastQC tool (0.11.9) [11], followed by removal of low quality bases (--nextseq-trim, Q<20), Illumina Universal adapter sequence and reads with less than 30bp length using Cutadapt(2.10) [12]. To access the quantity of host genomic DNA and other contaminants Kraken (2.0.9-beta) [13] was used and the reports were summarised using Krona(2.7.1)[14]. All the files were then aligned to human genome (assembly versionGRCh38) using HISAT2 (2.2.0)[15] and unmapped reads were extracted using SAMTOOLS(1.10)[16] and converted to FASTQ format using BEDTOOLS(2.29.2) [17] bamToFastq option.

### Alignment with viral genome

The unmapped reads were then aligned to SARS-CoV2 reference assembly (NCBI accession NC_045512) using HISAT2 (2.2.0) [15]. Amplicon primes from the aligned file were removed using iVar (1.2.2)[18] guided by Artic Network V3 primer scheme (https://github.com/artic-network/artic-ncov2019/tree/master/primer_schemes/nCoV-2019/V3). The aligned files were then de-duplicated using Picard Tools (2.18.7, https://broadinstitute.github.io/picard/). Alignment quality was checked using SAMTOOLS(1.10)[16] flagstat option.

### Consensus sequence generation and variant calling

Consensus sequence for each isolate was generated using Bcftools(1.10)[19] and SEQTK (https://github.com/lh3/seqtk). After generating a reference-based consensus sequence we selected 202 isolates with <5% N’s and more than 10x coverage for phylogenetic and mutation analysis. Single nucleotide variants were called and filtered (QUAL > 40 and DP >20) using Bcftools (1.10) [19]. Effect of the filtered variants were annotated using SnpEff (4.5)[20]. All of the consensus sequences were deposited in GISAID[21] (Supplementary table 1).

### Phylogenetic analysis

Phylogenetic tree of samples were performed using SARS-CoV-2 analysis protocol standards and tools provided by Nextstrain [22] pipeline. First all the sequences are aligned against the WH01 reference genome using Augur wrapper of MAFFT[23] and low quality variant sites are masked from the alignment. Initial tree was generated using Augur tree command using IQTREE algorithm[24]. The primary tree was further refinement of the tree was done using Augur refine command and the tree was rooted using the reference sequence with timeline information incorporation using TimeTree[25]. To finalize the tree for Nextstrain auspice visualization the tree was annotated using ancestral traits, clades, nucleotide mutation and amino acid mutation. The resulting tree was visualized using an Auspice instance.

### Haplotype network analysis

For haplotype network analysis we took a total of 287 (China 15, Germany 23, Italy 25, Saudi Arabia 23, Singapore 14 and South Korea) SARS-CoV-2 whole genome sequences with less than 1% N and with collection date of March, April and May from GISAID database. The selected samples are then aligned to the WH01 reference genome using MAFFT[23] After filtering aligned sequences we used POPART [26]software to generate haplotype network using Median Joining method with 2000 iterations.

### Modelling of the protein structures

The sequences of SARS-CoV2 proteins (NPS3, NSP4b, NSP6, Nucleocapsid, and Spike) were retrieved from NCBI. Since most of the proteins do not have a 3D structure in PDB, they were modelled using Modeller9.21 [27]. The suitable templates for modelling of the proteins (Supplementary Table 1) were selected by DELTA-BLAST[28] against the protein data bank (PDB) proteins. One hundred models were generated for each of the proteins. The best model was selected based on the lowest DOPE score [29]. The D614G mutant of the spike protein was generated by Modeller9.21. The loop in spike protein (670-690) was refined using loop modelling procedure in Modeller by generating 100 loop models. The model with the lowest dope score was finally chosen for both the mutant and wild type protein. Similarly, the host trans-membrane serine protease 2 (TMPRSS2) was also modelled using modeller. The PROCHECK [10] server was used to assess the stereochemistry of the generated models.

### Protein-Protein docking

The standalone version of HADDOCK2.2 [30] was used to perform protein-protein docking with SARS-CoV2 spike protein with human TMPRSS2. The docking was conducted by the restraining of the receptor (Spike) and ligand (TMPRSS2) residues known to be at the interface Spike-TMPRSS2 interface. Specifically, residues within 5 Å of the reported cleavage site (Arg685, Ser686) (Hoffman et al 2020) of the Spike and catalytic triad of TMPRSS2 (H296, D345, and S441) and binding residue D435 were restrained to be at the docking interface. A total of 100 docking poses were generated and ranked based on the HADDOCK2.2 docking score [30], which is composed of Van der Waals energy, electrostatic energy, restraints energy, etc. The best-ranked docking pose was visualized using Pymol.

### Statistical analysis and plotting

All the statistical analysis and plots were generated in R (3.6.1) statistical programming language using ggplot2, dplyr, reshape2, lubridate, ggsci and ggpubr package available from CRAN and Bioconductor (https://CRAN.R-project.org/package=tidyvers) repository.

## Acknowledgements

Authors would like to acknowledge Department of Biotechnology, India for providing the funding and approvals under the umbrella of DBT’s PAN-India 1000 SARS-CoV2 RNA genome sequencing consortium, ILS intramural grant for supporting for sequencing resources and Odisha state authorities to provide the clinical samples for testing from where the virus RNA were isolated for sequencing analysis. The authors do acknowledge GISAID, the genomic data sharing platform for influenza viruses and the contributors sequencing data used in this study. We gratefully acknowledge the efforts of volunteers who helped in co-ordinating the testing and related activities that supported implementation of this work.

## Author contributions

AP, SP, SR, SG, JT planned and designed the study; AJ, SM, MP, VKB, PSS, BS, NS, DS, AD, SK, AK, SS, JS did the experiments; SC, GHS, RD, SS, RS, TKB coordinated sampling and COVID-19 testing analysis; AG, SR, PP, did the genomic data analysis and SK and AD performed protein modelling and docking analysis. All authors have read and approved the manuscript.

## Supplementary Figure Legends

**Supplementary Figure 1: Subject demographics, transmission map and diversity of COVID-19 in India.**

(A-D) Represents the distribution of gender, age, clinical status and geographical location of our sampled subjects (n =225).

(E-F) Transmission map and Timetree of the 1042 high coverage Indian SARS-CoV2 whole genome sequences (N <1%) obtained from GISAID on 12th July, 2020.

**Supplementary Figure 2: ORF1ab Ct values in tested positive samples (n=637) and haplotype network of sequenced data coloured by clade.**

(A) ORF1ab Ct values in tested positive samples (n=637) binned in their respective collection months.

(B) Haplotype network of 202 SARS-CoV-2 whole genome sequences from our dataset coloured by their respective clade.

## Supplementary Table Legend

**Table S1:** List of single nucleotide mutations and the annotated genes, corresponding amino acid changes and its predicted impact on protein function.

