## Supplementary figures and images for "SARS-CoV2 genome analysis of Indian isolates and molecular modelling of D614G mutated spike protein with TMPRSS2 depicted its enhanced interaction and virus infectivity"

### Supplementary Figure 1

#### Supplementary Figure 1

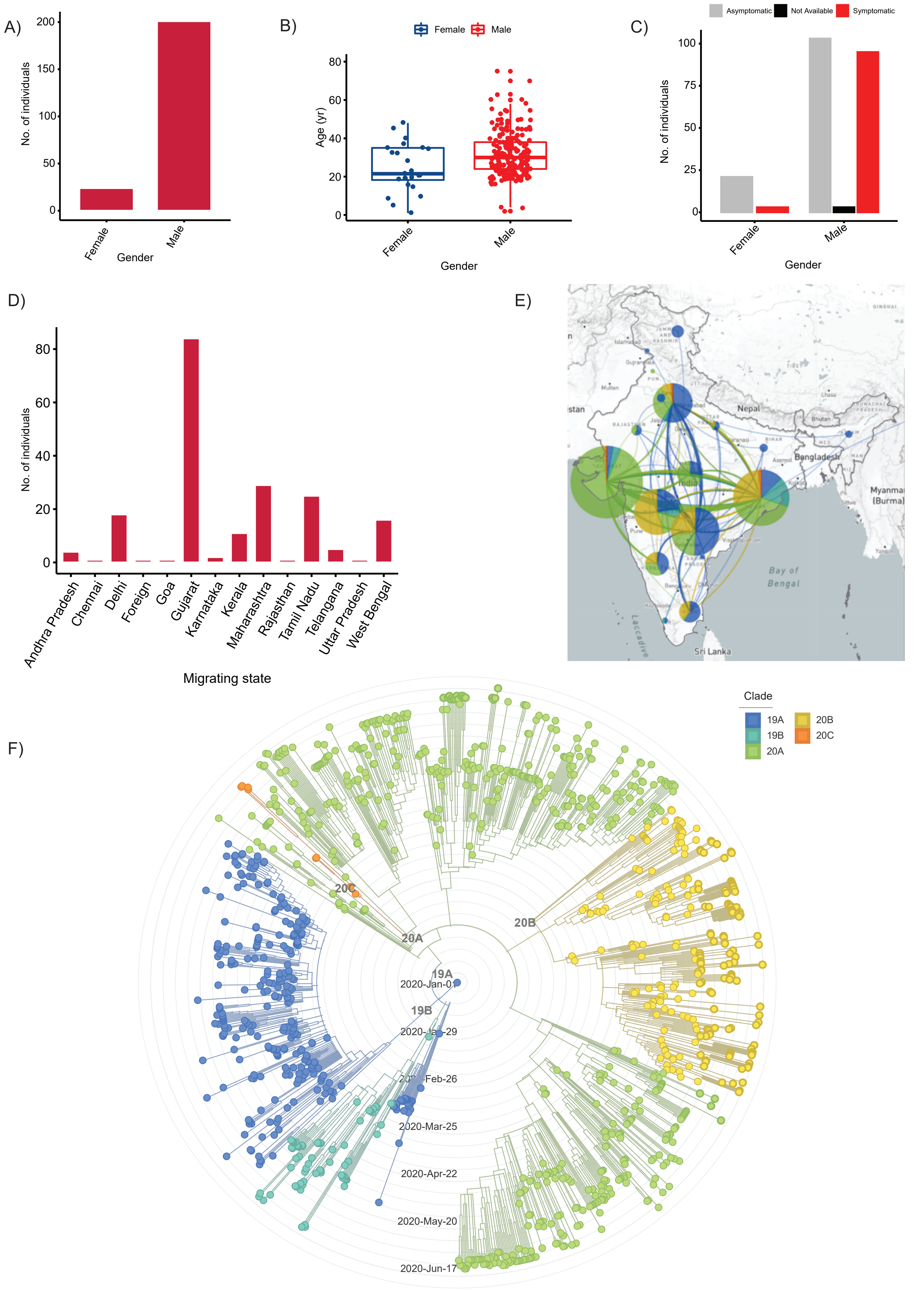

### Supplementary Figure 2

#### Supplementary Figure 2

A)

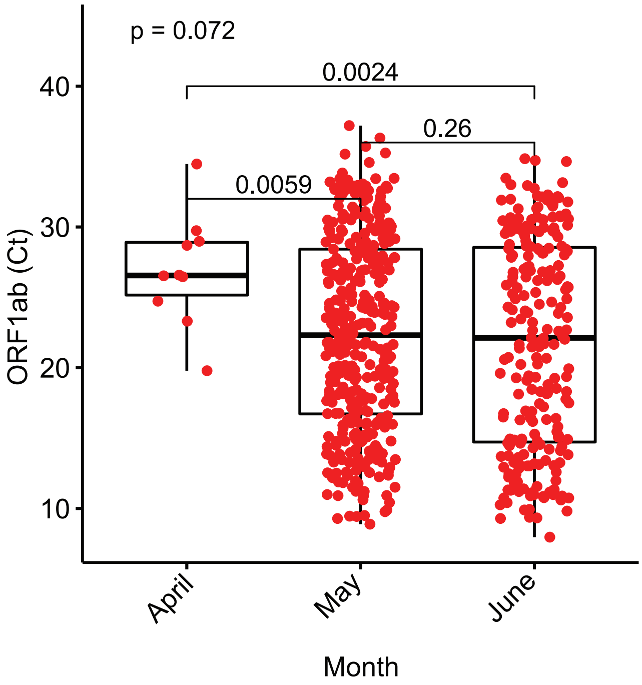


B)

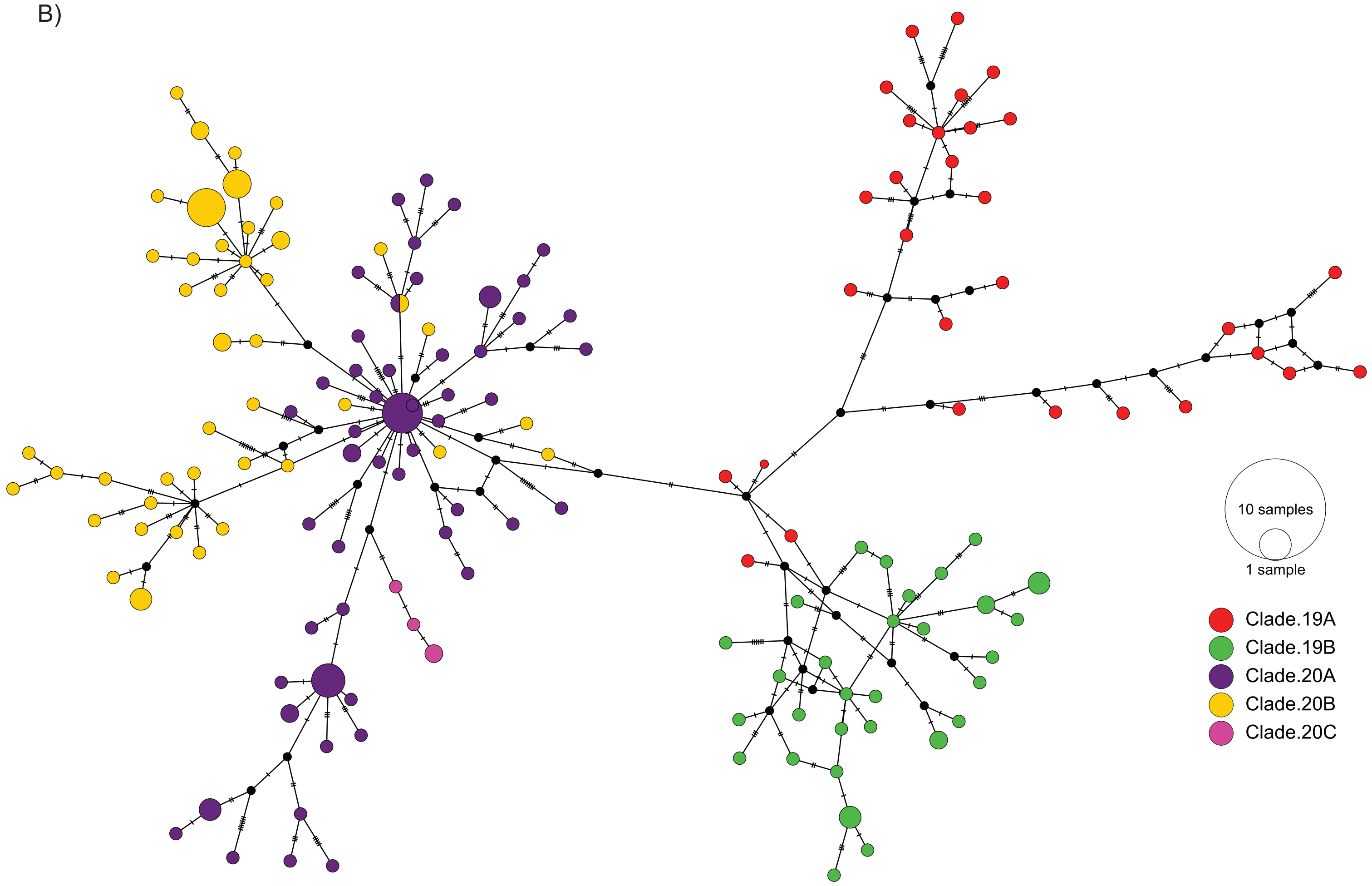
